## Supplementary figures and images for "Single-molecule analysis reveals the phosphorylation of FLS2 governs its spatiotemporal dynamics and immunity"

### Figure1-supplement-1

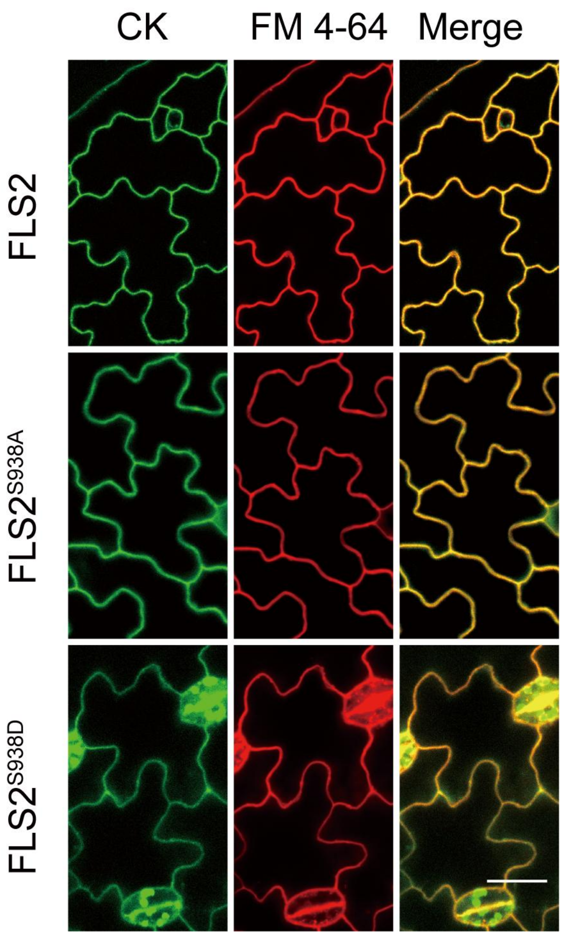

### Figure1-supplement-2

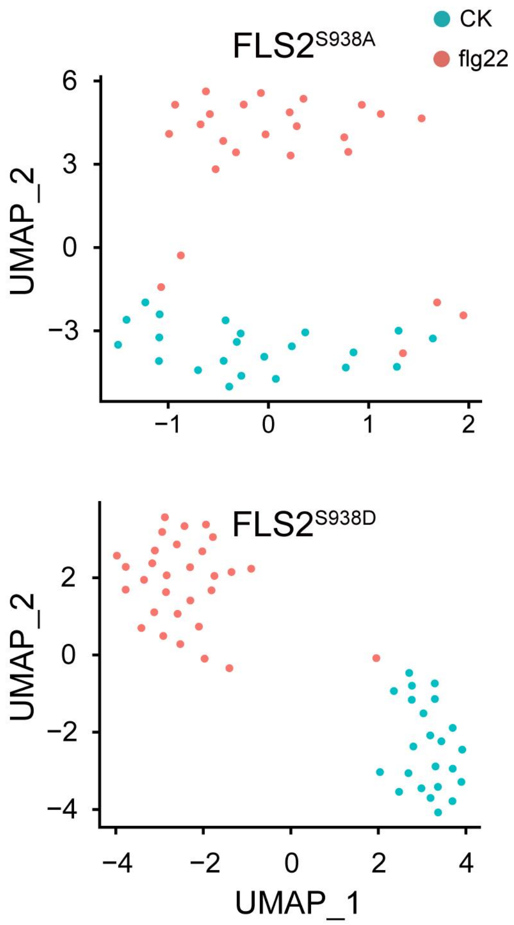

### Figure1-supplement-3

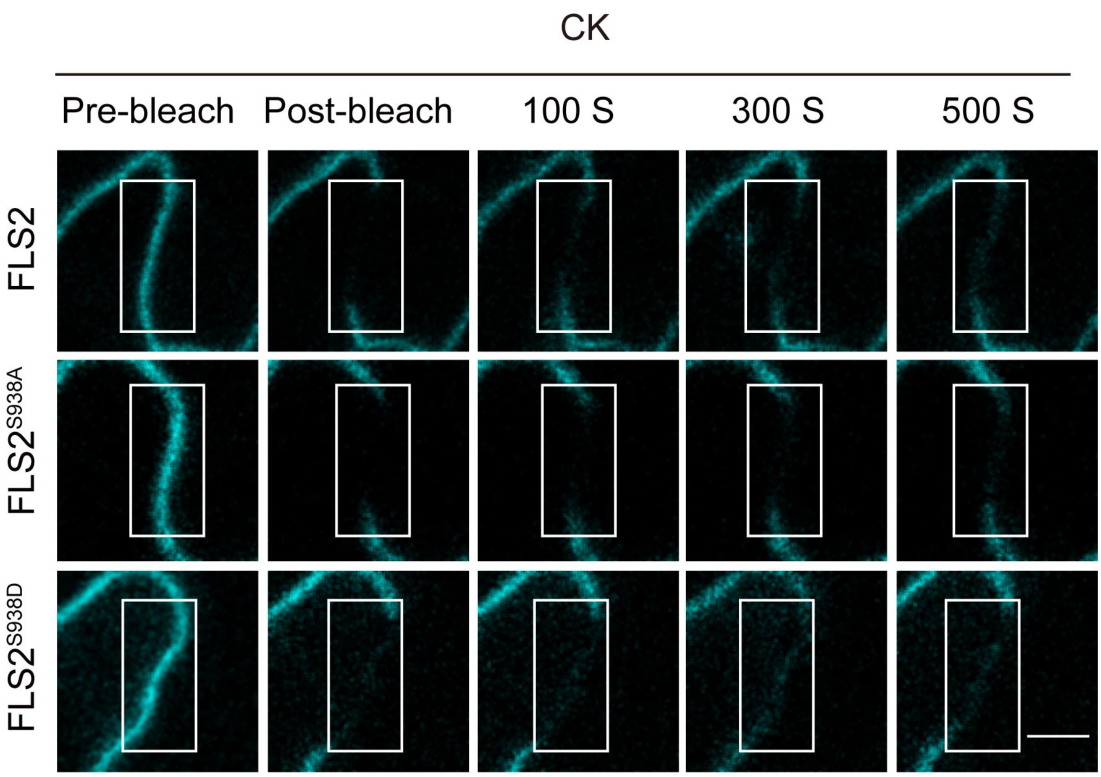

### Figure1-supplement-4

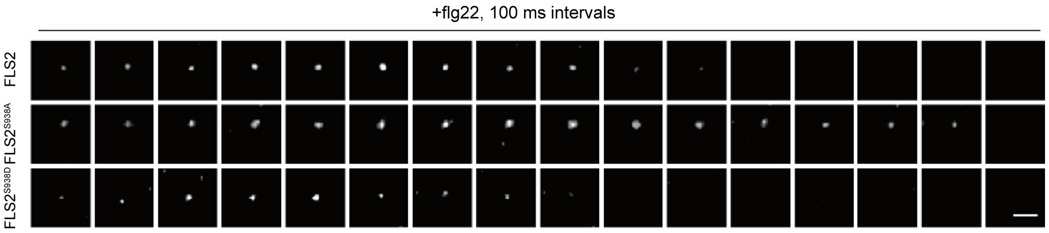

### Figure1-supplement-5

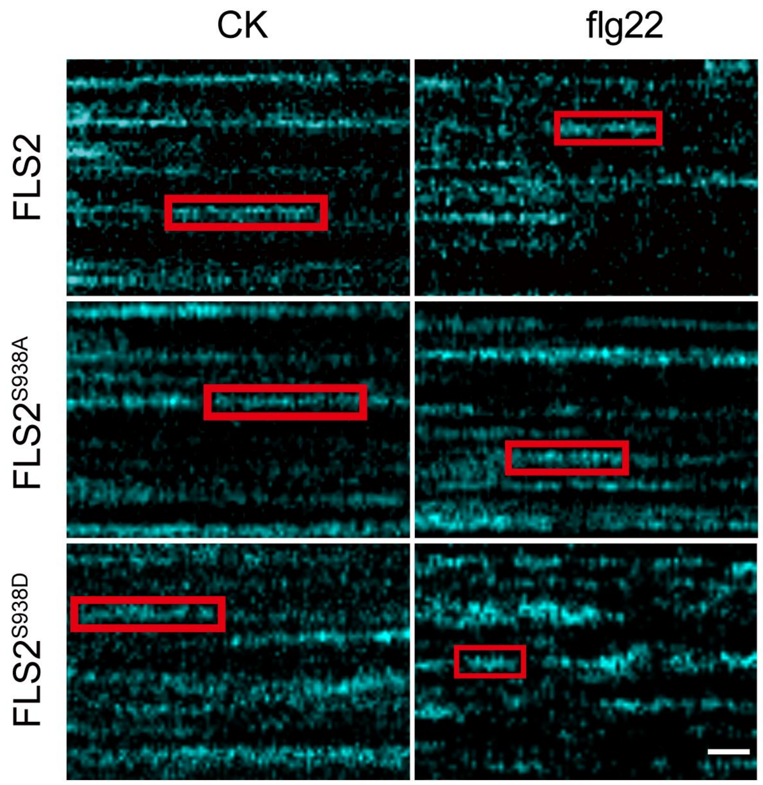

### Figure1-supplement-6

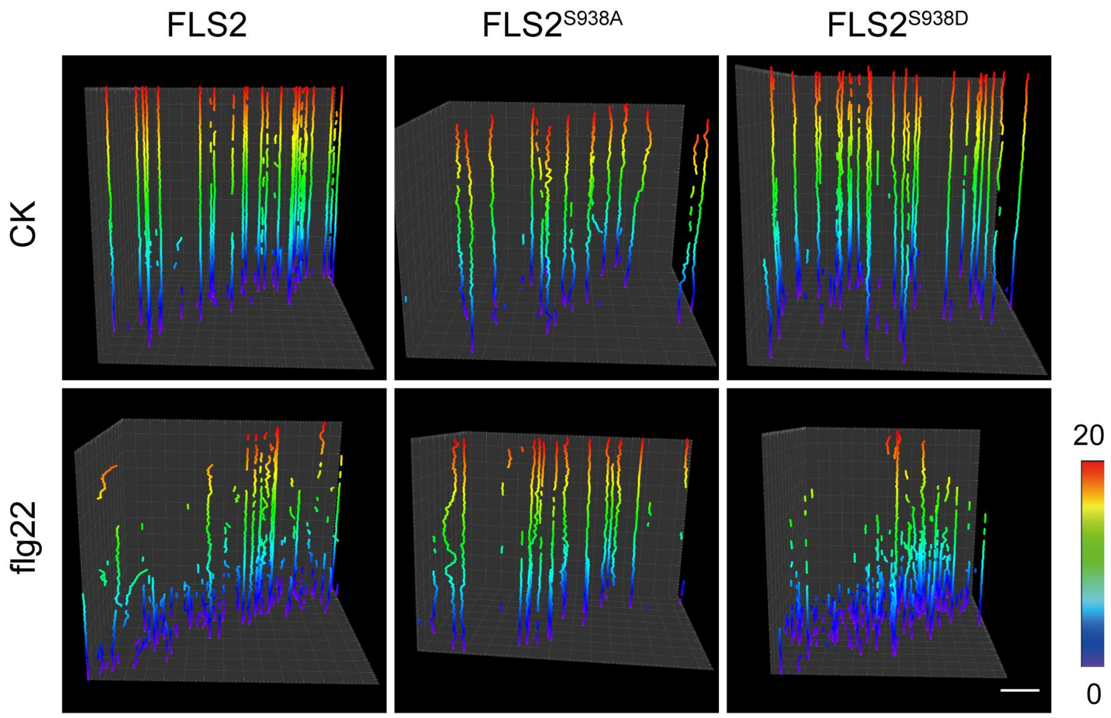

### Figure2-supplement-1

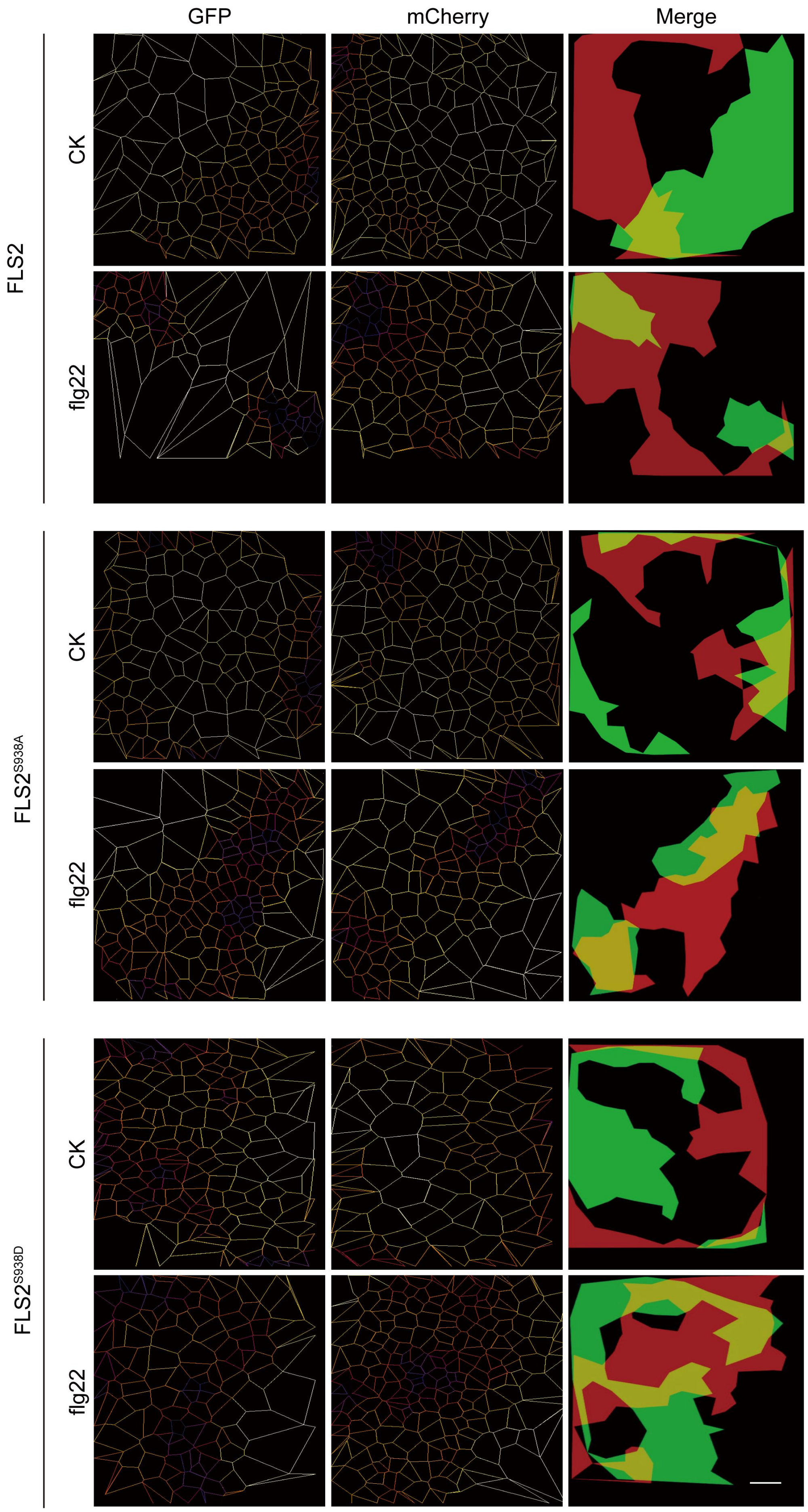

### Figure2-supplement-2

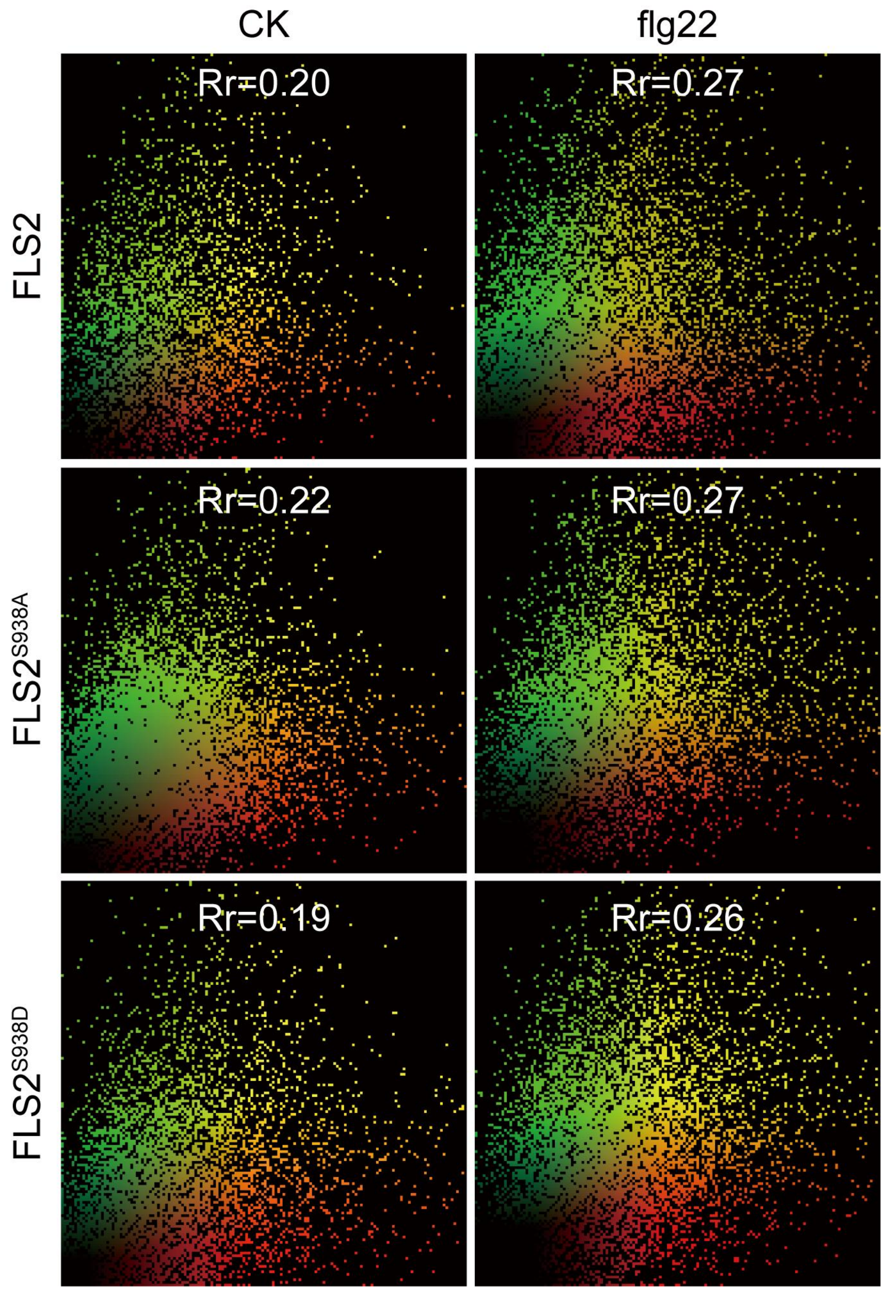

### Figure2-supplement-3

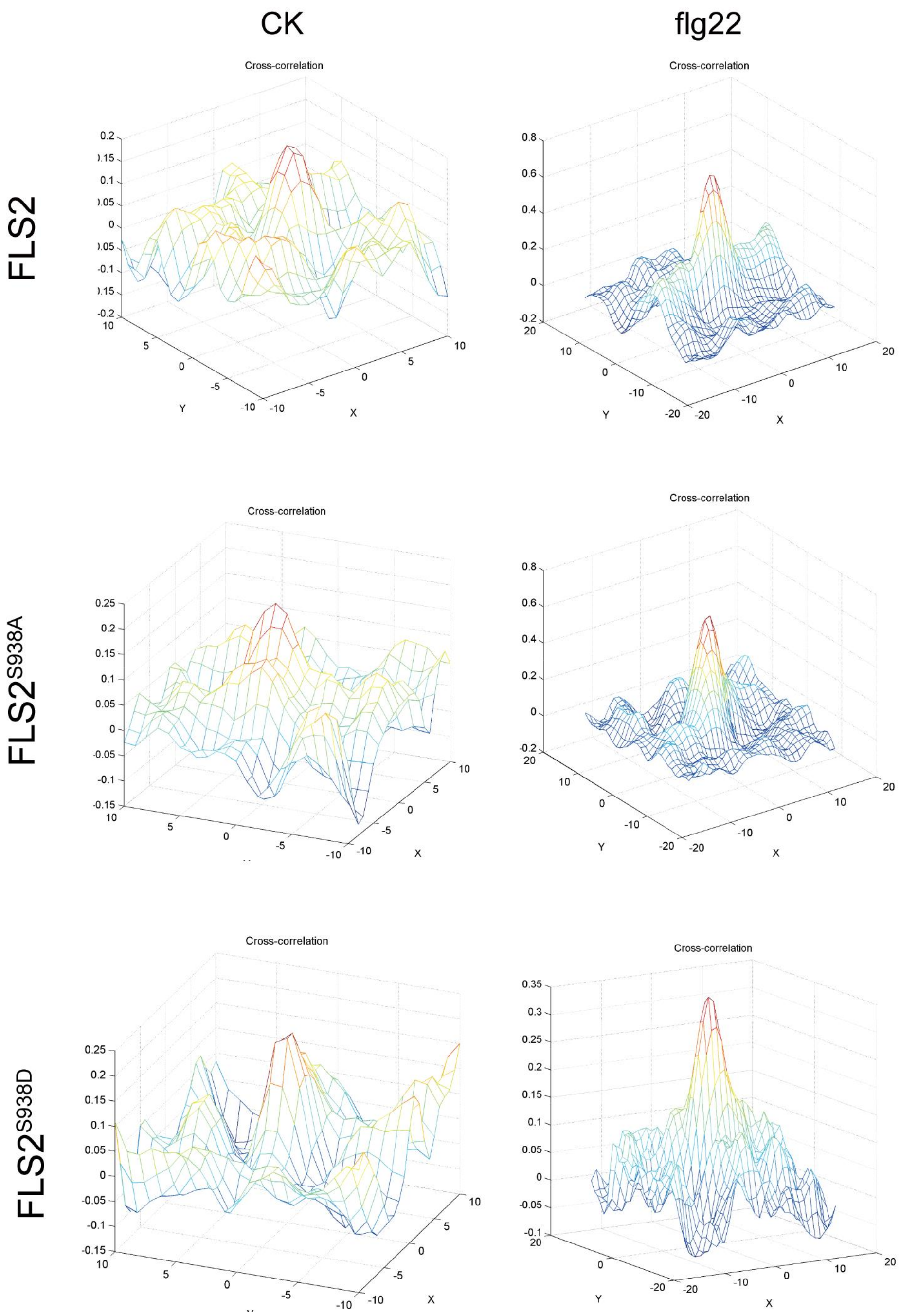

### Figure3-supplement-1

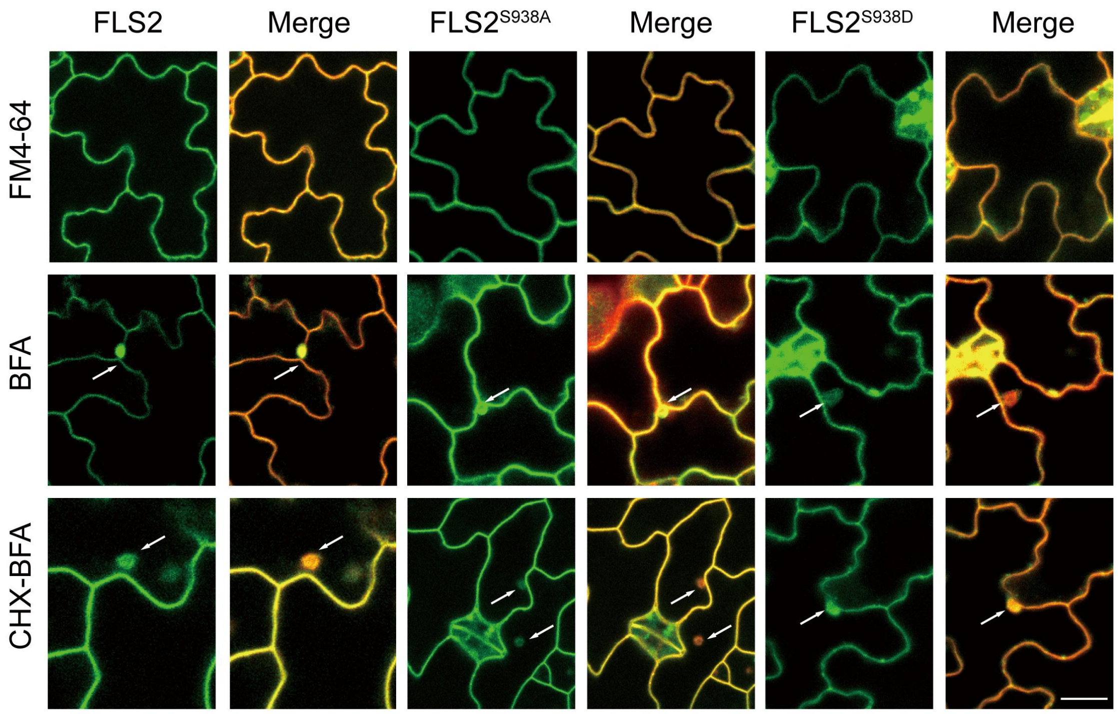

### Figure3-supplement-2

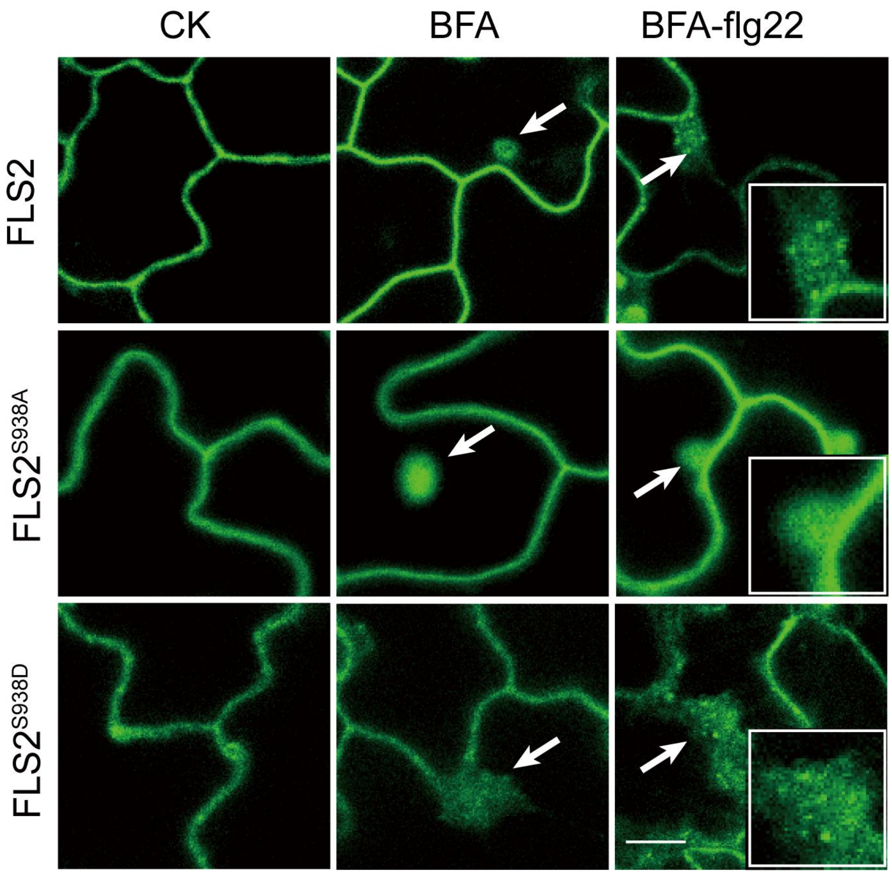

### Figure3-supplement-3

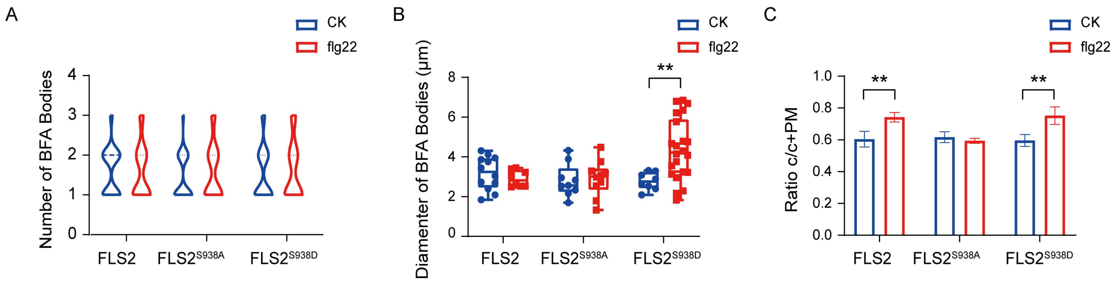

### Figure3-supplement-4

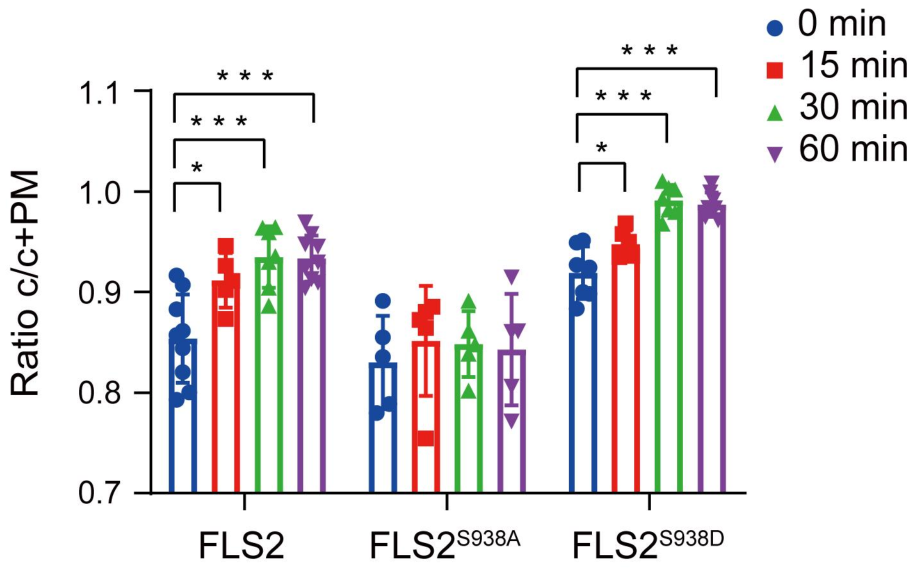

### Figure3-supplement-5

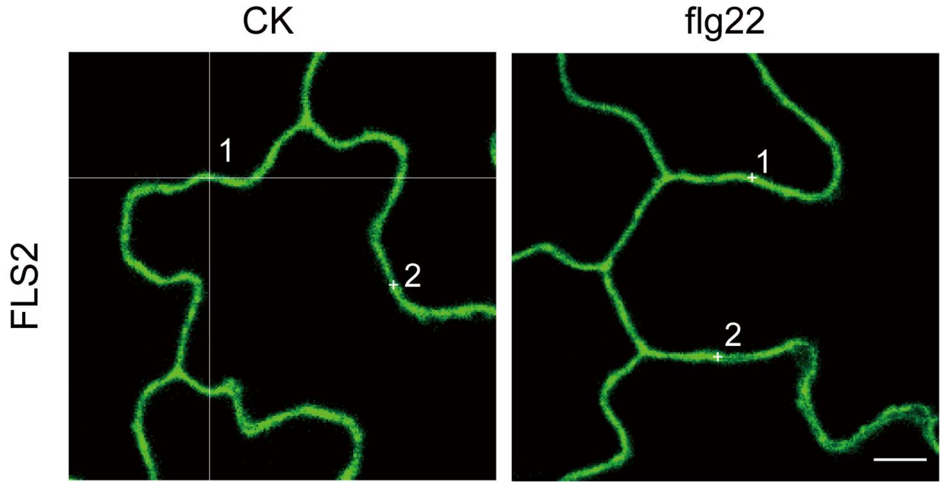

### Figure4-supplement-1

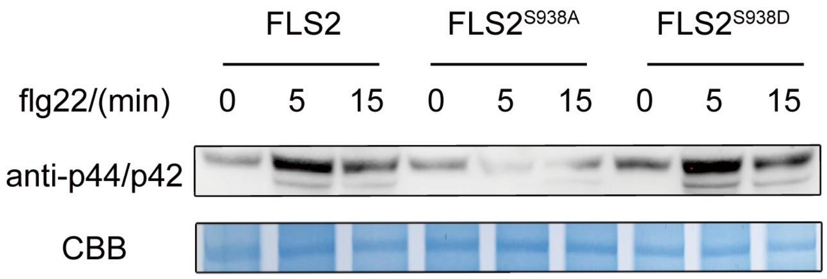

### Figure4-supplement-1

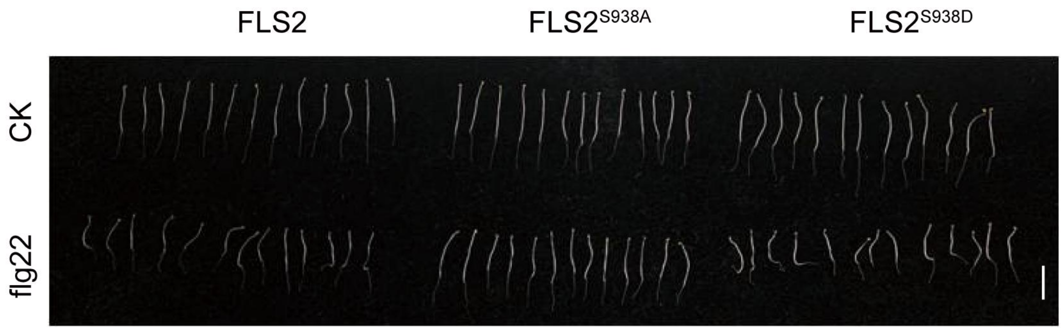

### Figure4-supplement-2

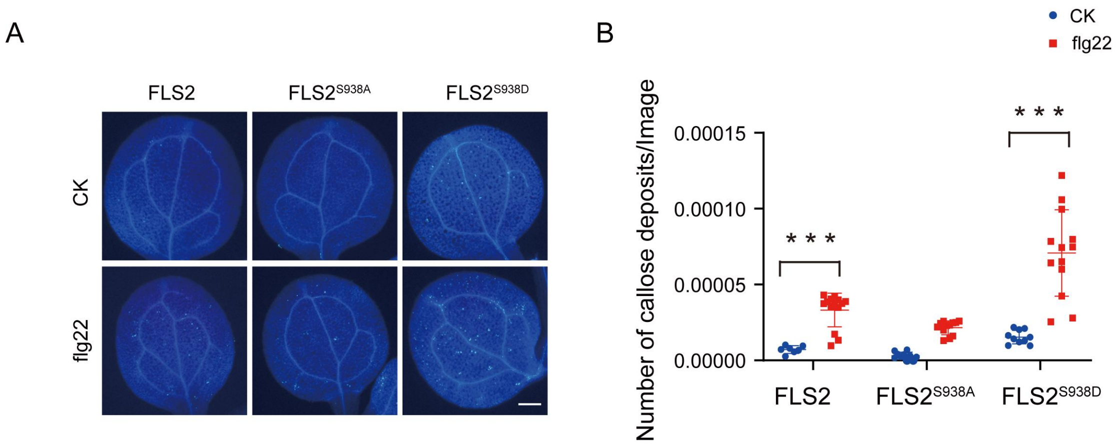
